# Experimental investigation of temperature-dependent denaturation behavior of type-I Collagen

**DOI:** 10.1101/682401

**Authors:** İ. Deniz Derman, Esat C. Şenel, Onur Ferhanoğlu, İnci Çilesiz

## Abstract

Precise investigation of the temperature and the duration for collagen denaturation is critical for a number of applications, such as adjustment of temperature and duration during a laser-assisted tissue welding or collagen-based tissue repair products (films, implants, cross-linkers) preparation procedures. The result of such studies can serve as a guideline to mitigate potential side effects while maintaining the functionality of the collagen. Though a variety of collagen denaturation temperatures have been reported, there has not been a systematic study to report temperature-dependent denaturation rates. In this study, we perform a set of experiments on type-I collagen fiber bundles, extracted from the rat-tail tendon, and provide an Arrhenius model based on the acquired data. The tendons are introduced to buffer solutions having different temperatures, while monitoring the contrast in the crimp sights with a wide field microscope, where collagen fibers bend with respect to their original orientation. For all tested temperatures of 50°C–70 °C and tissues that were extracted from 5 rats, increasing the temperature reduced the contrast. On the average, we observed a decay of the contrast to half of its initial value at 37, 157, and 266 seconds when the collagen was introduced to 70 °C, 65 °C, and 60 °C buffer solutions, respectively. For the lower temperatures tested we only observed to be only about 20% and 2 % decay in the crimp contrast after > 2 hours at 55 °C and 50 °C, respectively. The observed denaturation behavior is also in line with Arrhenius Law, as expected. We are looking forward to expand this study to other types of collagen as a future work. Overall, with further development the data and model we present here could potentially serve as a guideline to determine limits for welding and manufacturing process of collagen-based tissue repair agents.

## 1 Introduction

A great number of literary studies have focused on characterizing temperature dependent properties (mechanical, electrical, optical, etc.) of collagen or collagen-rich tissue. For instance Chae et al [1] have studied mechanical changes of cartilage tissue (composed of 3D fibrillar collagen networks) under RF or contact heating treatment, which is critical in understanding the permanent effects caused on the tissue site of interest. Collagen and gelatin systems were previously observed under different temperatures with a dilatometer setup to identify phase transitions [2]. Calorimetric investigations were conducted on collagen tissue to reveal collagen denaturation temperature [3,4], where some studies focused on age-related changes in denaturation behavior [5]. High-resolution imaging techniques (scanning electron microscopy [6,7], optical coherence tomography [8]) as well as spectroscopy (Fourier Transform Infrared [3] and Fluorescence spectroscopy [9]) have also been widely utilized to observe denaturation behavior. In these studies, the reported denaturation temperatures vary within a wide range of 45 – 70 °C.

Here, we present the results and Arrhenius modeling based on a systematic set of experiments conducted on type-I collagen, observing their denaturation rates and durations at different temperatures. The denaturation rate vs. temperature behavior is crucial for a number of biological and biomedical applications. Laser assisted tissue welding [10,11] is one application we want to stress out, where performing a welding procedure under denaturation temperature and duration will allow for minimal tissue damage while bonding disjoint tissue sites.

One other application that our study could be beneficial is adjustment of collagen-based tissue bonding and repair products such as polymeric hydrogels [12], titanium implants coated with collagen (or alike proteins) and collagen scaffolds for bone regeneration [13,14]. Likewise tissue welding application, care should be taken in manufacturing of such medical products in the any heat-involved steps to ensure collagen is still functional at the end.

## 2 Methods

The experiments were conducted on Wistar Rat (n = 5, ~ 10 months old, weight: 380 – 400 gr) tails, which were received from and dissected at the Central Vivarium of Boğaziçi University. The dermal layers were first removed from the tail to expose the skeletal frame and the collagen-I rich tendons. Then, tendons were lifted from the skeletal structure and stored in 50 ml phosphate buffered saline (PBS). For each experiment, a single tendon was isolated and sandwiched between two glass slides to be inserted in a heat controlled PBS solution, as illustrated in Fig 1a.

**Figure 1.**
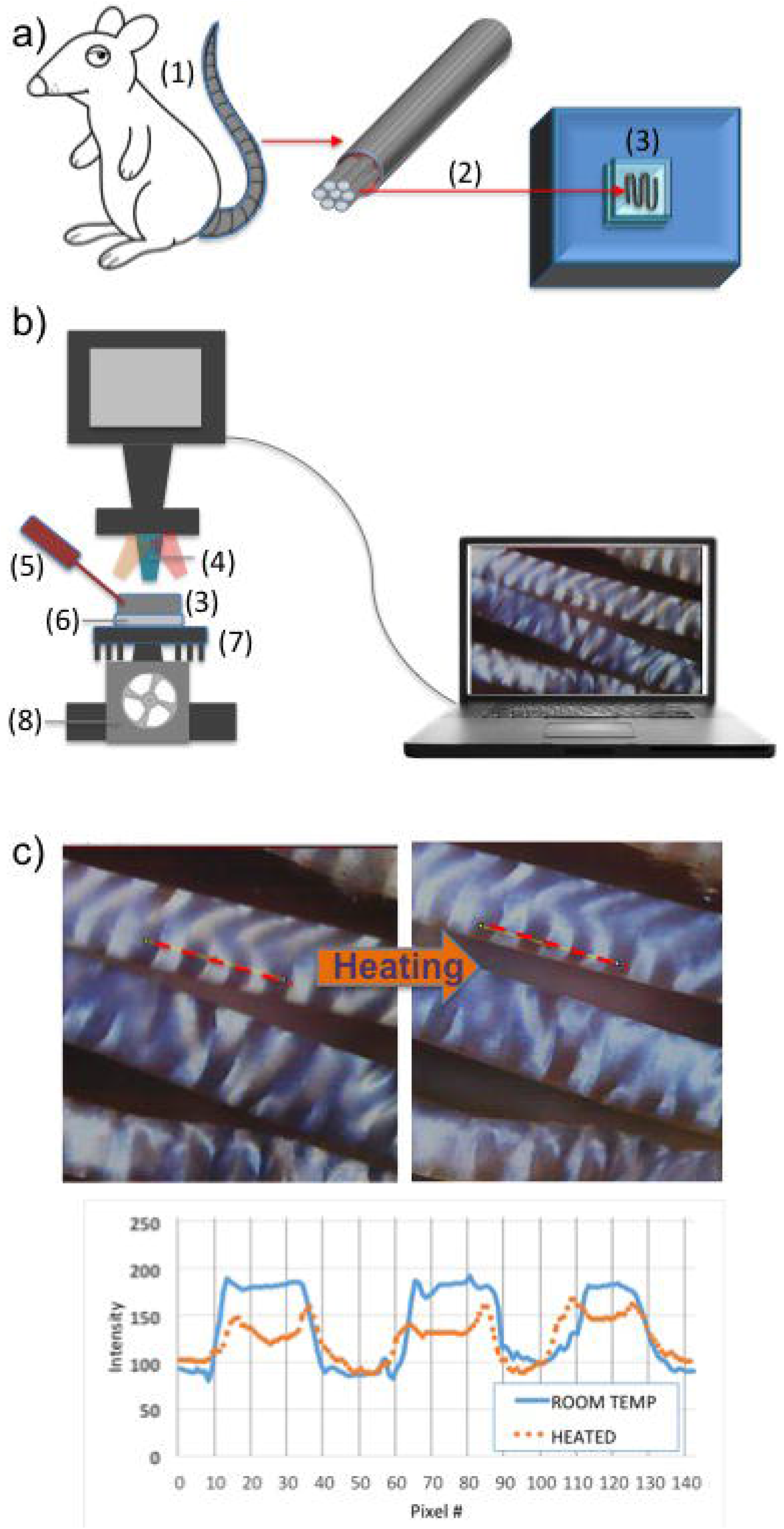
**a)** preparation of tissue samples from Wistar rat (1) tail (2) and placement into the sample cuvette (3). **b)** Microscopy setup involved in the experiments, comprising of an objective lens (4), temperature sensor (5), TEC (6), aluminum fins (7), fan (8). **c)** An exemplary screen-shot from the microscope and plot of the highlighted crosssection for nominal temperature and heated tissue shows the decrease in crimp contrast.

The extracted tendon was twisted in a meander fashion right before it was placed between two glass slides, to maximize the amount of tissue that can be visualized with the microscope setup. An upright microscope, illustrated in Fib 1b was employed to monitor collagen fibers. A thermoelectric-cooler (TEC) was placed under the aluminum sample cuvette and is utilized to control the PBS temperature in which the tendon resides, while monitoring the temperature with a thermo-couple unit that is connected to an Arduino platform. Metal fins and a fan was also incorporated into the setup to better distribute the heat extracted from the bottom side of the TEC. A total of 5 rat-tails and 2 sites of each tail were video observed under 50 °C, 55 °C, 60 °C, 65 °C, and 70 °C, for a maximum duration of ~ 8 hours (for 50 °C).

Under the microscope, collagen fibers appear crimped, with alternating dark and bright regions that are nearly orthogonal to the fiber direction [15]. Previous research has shown that the crimps, observed via scanning electron microscopy (SEM), are sites where collagen fibrils suddenly change directions [16,17]. Changes in the crimp patterns and structures have been monitored investigating the effects of strain [18], and rupture [19] on collagen tendons. In this study, we propose to use crimp contrast as an indicator of denaturation.

The contrast of the crimp patterns improve upon using a polarized microscope, yet the crimps are also observable under a regular wide-field microscope, as shown in Fig 1c, and also with the bare eye. Here, we define crimp contrast (*γ*) as:

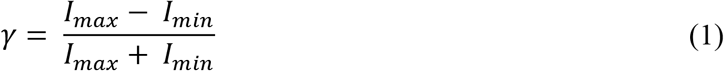

where *I_max_ is* the average intensity of the selected bright stripe and *I_min_* is the average intensity of the adjacent dark stripe of the collagen crimp. In our experiments, the crimp contrast was time-lapse monitored for all 5-rat tails for all listed temperatures, and 2 chosen locations per tail. Fig 1c illustrates a tendon, and its intensity for the dashed cross-section. Note that temperature increase results in a diminished crimp contrast.

## 3 Results

Figure 2a illustrates the change in the normalized crimp contrast (*γ*) on five rat-tails, two locations per tail, observed at at 5 different temperatures. As expected the decay in contrast is more rapid at high temperatures, and significantly slows down at lower temperatures. Finally, we observe no significant signs of contrast change at 50 °C for a total duration of 30,000 seconds (> 8 hours).

**Figure 2.**
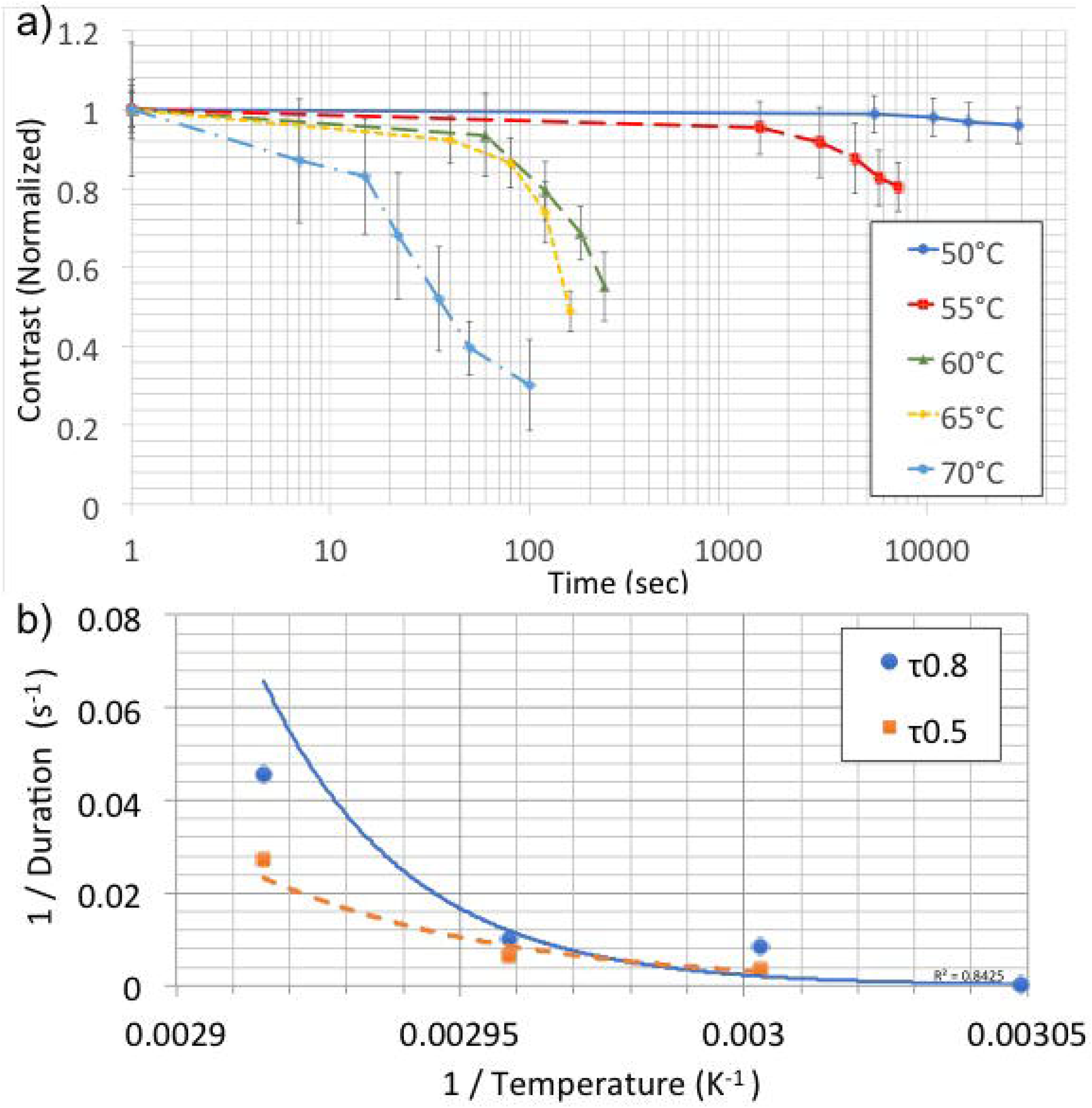
**a)** Decay in crimp contrast (normalized to initial value) as a function of time, at different buffers solution temperatures. **b)** Arrhenius plots of average rates (inverse durations: τ_0.5_^−1^ and τ_0.8_^−1^) with respect to temperature. Dashed and solid lines are exponential fits to the shown data values.

The error bars on the figure reflect the standard deviation of the contrast, for 10 different data from 5 different tissues. For any temperature, we identify the average duration needed for the contrast to drop down to 50% of its initial value, τ_0.5_, as the duration for denaturation, and τ_0.5_^−1^ as the denaturation rate. Note that experimental results reveal an average denaturation duration of τ_0.5_ = 37, 157, 266 seconds for 70 °C, 65 °C and 60 °C, respectively. For lower temperatures the denaturation duration was not reached within the timeframe of the experiments. We also define the duration needed for the contrast to drop down to 80% of its initial value, τ_0.8_, as the duration for onset of denaturation, and τ_0.8_^−1^, as the rate for onset of denaturation. Once again, the experimental results reveal an average duration for onset of denaturation as τ_0.8_ = 22, 100, 120, and 7200 seconds for 70 °C, 65 °C, 60 °C and 55 °C, respectively.

Without loss of generality, the decay of the contrast to other values than 50% or 80 *%* of its initial value could also be chosen, given that any guidelines for aforementioned biomedical applications should impose smaller durations (by at-least a few folds) than that of the duration for denaturation that the collagen experiences at any temperature.

Arrhenius plot is a simple model and has been widely used in describing thermal denaturation of various proteins and cells [20]. Thus, as a next step we model the acquired crimp contrast rates based on Arrhenius equation that links reaction rate to inverse temperature, such that:

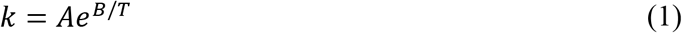

where *k* is the rate constant, taken as either τ_0.5_^−1^ or τ_0.8_^−1^, *A* is the pre-exponential factor, *B* is the exponential factor (activation energy) that determines the rate, and *T* is temperature in Kelvins. Plotting τ_0.5_^−1^ or τ_0.8_^−1^ as a function of inverse temperature (Kelvins), we observe the plots in Fig. 2b, where we observe an R^2^ value of 0.84 and 0.94 for τ_0.5_^−1^ and τ_0.8_^−1^ data with the exponential fit lines.

## 4 Conclusions and Discussion

In this study, through a systematic set of experiments we observed the decay in crimp contrast of rat tail tendons (n=5) at different temperatures, as a function of time,. The experiments revealed that a decay of crimp contrast to half of its initial value within 37, 157, 266 seconds, for tested temperatures of 70 °C, 65 °C and 60 °C, respectively. The experiments conducted at 55 °C and 50 °C showed a decay in crimp contrast of < 20 %. The obtained results were also observed to match the traditional Arrhenius model well, which has priorly been utilized in cell and protein denaturation studies. Investigation of denaturation rates as opposed to the denaturation temperature, allows for providing guidelines for a number of biomedical applications, such as adjustment of temperature and duration during a laser-assisted tissue welding, or the heat treatment steps while manufacturing collagen-based tissue repair products (films, implants, cross-linkers). As a future work, we are looking forward to expand this study to other types of collagen towards forming a guideline for biological / medical procedures that require heat in order to mitigate their side effects of denaturation of collagen or other proteins.

## Acknowledgments

We acknowledge Boğaziçi University’s Central Vivarium for their assistance in the preparation of rat tail tissue.

---

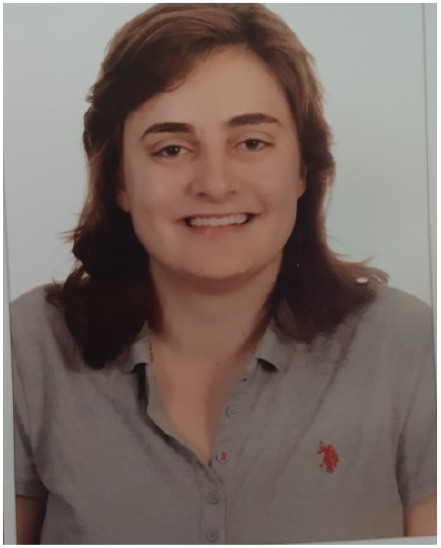

**İ Deniz Derman** received the B.S. degree in Biomedical Engineering from Bahçeşehir University in 2018. Currently she is an M.Sc. student of Biomedical Engineering and a research assistant of the Electro-Optical Devices Laboratory in Istanbul Technical University. Her research interests are 3-dimensional optomedical imaging, collagen extraction and optical characterization of proteins. She is the recipient of a number of awards, including P&G project award and the IEEE Darrel Chong Student Activity Award.

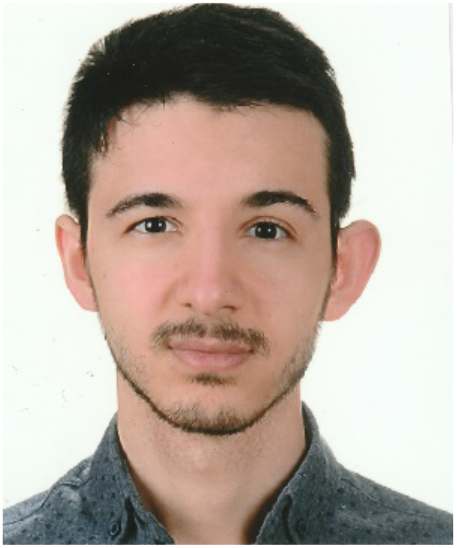

**Esat C. Şenel** received the B.S. degree in Biomedical Engineering from Bahçeşehir University in 2018. Currently He is an M.Sc. student at Istanbul Technical University. His research interests are microfluidics and optical characterization of proteins.

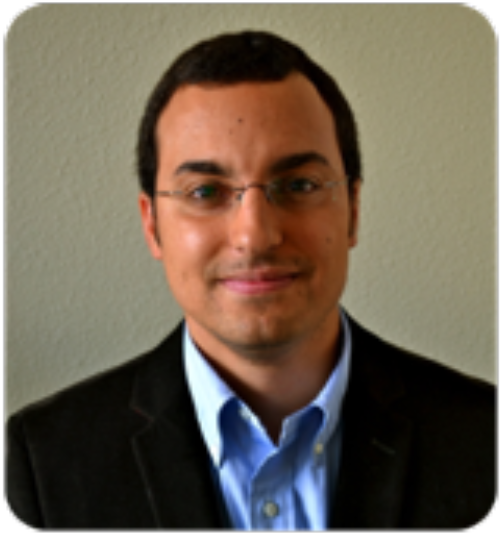

**Onur Ferhanoğlu** received B.S. and M.S. degrees from Bilkent University, in 2003 and 2005 respectively, in Electrical Engineering. In 2005, he joined the Optical Microsystems Laboratory at Koç University as a graduate researcher, where he developed MEMS based thermal imaging sensor arrays. During graduate studies, he visited Johns Hopkins University (2004), Georgia Tech. (2007) and EPFL (2010) as a research scholar. After receiving his Ph.D. (2011), he became a post-doctoral fellow at Femtosecond Laser Assisted Biophotonics Laboratory at the University of Texas at Austin, where he played a key role in the development of an ultrafast laser microsurgery scalpel (2011-2014). Dr. Ferhanoğlu is now appointed with the Electronics and Communication Engineering department of Istanbul Technical University. His research interests include biomedical optics, and MEMS for medical applications.

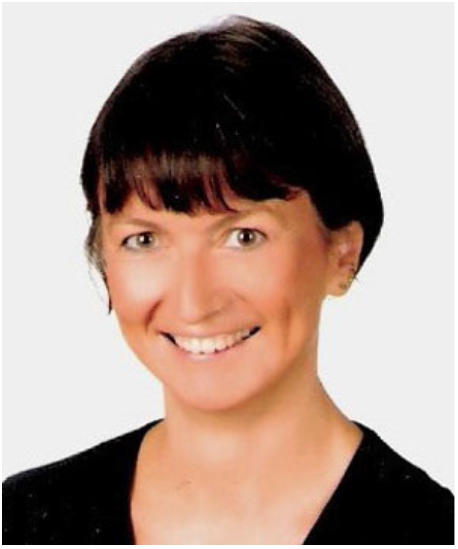

**İnci Çilesiz** received the B.SE degree in Electronics and Communication Engineering from İstanbul Technical University in 1982. She then joined the Biomedical Engineering department of University of Texas at Austin, where she focused on tissue optics and laser-assisted tissue welding (1989-1994). Since then, she has been a faculty with the Electronics and Communication Engineering of Istanbul Technical University, where she is currently a full professor. She has been a visiting researcher at University of Amsterdam (2000-2002), Tel-Aviv University (1998), and University of Minnesota (2007). Her research interests fall within the broad field of biomedical engineering, with a focus on biomedical instrumentation, temperature dependent optical properties of tissues, and biomedical optics.

